# Effect of accelerated ageing on seed membrane integrity and chemical composition of *Tetrapleura tetraptera* (schum. & thonn.)

**DOI:** 10.1101/662122

**Authors:** H. S. Sossou, J. M. Asomaning, E. A. Gaveh, A. F. K. Sodedji, E. E. Agoyi, J. Sarkodie-Addo, A. E. Assogbadjo

**Affiliations:** CSIR-Forestry Research Institute of Ghana, Fumesua, Kumasi-Ghana; Department of Horticulture, Kwame Nkrumah University of Science and Technology, Kumasi, Ghana; Laboratory of Applied Ecology, Faculty of Agronomic Sciences, University of Abomey-Calavi, Cotonou, Benin; Department of Crop and Soil Sciences, Kwame Nkrumah University of Science and Technology, Kumasi, Ghana

**Keywords:** Seed ageing, Membrane integrity, Chemical composition, Storage potential

## Abstract

Seed ageing is one of the major issues in the storage of seeds. Accelerated ageing test has shown great potential for studying the mechanism of ageing and associated deterioration processes of seeds. This study was conducted to evaluate the effects of accelerated ageing process on seed membrane integrity and chemical composition of Tetrapleura *tetraptera* (schum. & thonn.). Seeds were subjected to traditional accelerated ageing (100% RH at 38 and 41°C for 48, 72 and 96 h) and salt-saturated accelerated ageing (76% RH at 38 and 41°C for 48, 72 and 96 h) tests and the aged seeds were tested for moisture content, leachate conductivity and chemical composition. Results showed that the accelerated ageing process led to an increase in seed moisture content, carbohydrate, fat, ash and crude fibre content and a reduction in crude protein content in the aged seeds. Also, the leachate conductivity test revealed a loss in membrane integrity of T. *tetraptera* seed indicating that the accelerated ageing process triggered the seed coat dormancy release in T. *tetraptera* seed and led to the leaching of the mineral constituent of the seed. The accelerated ageing test combined with a storage experiment could therefore be a promising technique for estimating the storage potential of T *tetraptera* seed but further investigations including the enzyme activity, are required to standardize the appropriate testing conditions for the species.

## INTRODUCTION

Seed vigour testing is very crucial for sustainable conservation of forest tree species. Various factors reduce the vigour of stored seeds resulting in low germination and poor field establishment. The commonly known factors which influence vigour level include: genetic; environmental conditions; seed maturity at harvest, physical characteristics, mechanical integrity, deterioration and ageing, and pathogens. Recent studies have focused on the physiological causes of differences in seed vigour, mainly the role of seed ageing (Copeland and McDonald, 1995). Seed ageing as a result of a physiological deterioration generally starts at physiological maturity and continues during harvest, conditioning and storage (Kapoor et al., 2011). The deterioration process in seed can be described as cumulative, irreversible, degenerative and inexorable (Kapoor et al., 2011) and is accompanied with numerous cellular, metabolic and chemical alterations as well as chromosome aberrations and damage to the DNA, impairment of RNA and protein synthesis, modifications in the enzymes and food reserves and loss of membrane integrity (Kibinza et al., 2006). Deterioration can occur in seed in a few days or years but becomes evident as germination percentage reduces and produces weak seedlings, leading ultimately to its death. However, its rate depends on several factors including, seed moisture, storage temperature, and genetic factors (Walters et al., 2010).

Accelerated ageing test has shown great potential for studying the mechanism of ageing and associated deterioration processes of seeds. It is generally done by subjecting seeds to high temperature and high relative humidity to increase the rate at which the various changes occur within the seed and provide in a relatively short period of time, indices for estimating the physiological potentials of seed lots. Hence, the accelerated ageing techniques have received extensive testing for their efficiency as tree seed vigour test (Bonner, 1998).

*Tetrapleura tetraptera* (Schum and Thonn), is an important indigenous tropical tree species of Fabaceae family. It is distributed across the rainforest belts of West, Central and East Africa. The species exhibits an undeniable socio-economic importance for the local communities across its distribution area as result of its numerous nutritional and medicinal properties (Adewunmi et al., 1991). In spite of that, the natural population of *T. tetraptera* is declining at an alarming rate due to its over-exploitation (Nya et al., 2000). Like any other forest tree, *T. tetraptera* regeneration cycle is long and the natural germination of the seeds is still a challenge for its establishment (Jimoh, 2005). Further, *T. tetraptera* exhibits poor germination when freshly collected (Nya et al., 2000). These intrinsic factors make the species not quite adequate for use in reforestation programmes. To promote its ex-situ conservation and ensure timely availability of quality seed for forest rehabilitation programmes, Sossou et al., (2017) investigated different seed testing techniques on the species, including accelerated ageing test. Till date, the physiological and chemical changes that occurred in the seeds under the different tested conditions were not documented. This study aims to determine the effect of different ageing conditions on seed membrane integrity, and chemical composition of *T. tetraptera* seeds in order to trace the deterioration process in aged seed and provide valuable information for predicting the storability of *T. tetraptera* seed lots using accelerated aging test.

## MATERIALS AND METHODS

This study was carried out in the laboratory of the National Tree Seed Centre located at the CSIR-Forestry Research Institute of Ghana in Kumasi and the Laboratory of Horticulture, Faculty of Agriculture and Natural Resources, Kwame Nkrumah University of Science and Technology, Kumasi. The experimental materials include eight seed lots of *T. tetraptera* obtained from the National Tree Seed Centre of the Forestry Research Institute of Ghana in October 2015. The seeds were cleaned, kept for a month at ambient temperature (24-28°C) and subsequently used for the various experiments.

### Accelerated ageing

Two different tests were performed in the ageing experiment: the traditional accelerated ageing and salt-saturated accelerated aging. For the traditional accelerated ageing (TAA) test, seeds were aged at 100% relative humidity (RH) using distilled water (ISTA, 1987). The salt-saturated accelerated ageing (SSAA) was carried out using the procedure proposed by Jianhua and McDonald (1996), which consists in replacing the distilled water with the same amount of a saturated NaCl solution (obtaining 76% RH). Both tests were conducted in clear plastic containers (boxes) having suspended wire mesh screen inside, in which 80 g of seeds were spread to form a single layer. The screen was positioned in the plastic containers 6 cm above the water level and the boxes covered with lids. All samples were placed in an incubator at 38 and 41°C for 48, 72 and 96 h.

### Solute leachate test

This test was carried out to assess the effect of ageing procedures on the *Tetrapleura tetraptera* seed membrane. Four replicates of twenty seeds were taken from each sample and weighed in grams to two decimal places prior to testing. Each replicate was rinsed twice with 20 ml of deionized water and placed in 150 ml beaker, stirred and covered to avoid pollution and evaporation of water. Two beakers with the same quantity of de-ionized water were set up as blank. All beakers were kept on laboratory bench at the room temperature (24-28°C) at which the cell was calibrated. The conductivity of the leachate solution was measured and recorded after 1, 2, 3, 4, 5, 6 and 24 hours. The conductivity dip cell was rinsed once between each sample measurement. Specific conductivity per grams of dry seed was calculated using the following formula (ISTA, 1987):

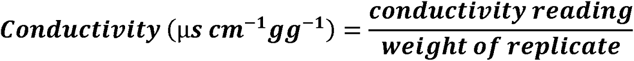

### Moisture content determination

This was done by using the low constant temperature (103±2 °C for 17h±1) gravimetric method as described by ISTA 2007.

### Chemical composition determination

Ash, crude fibre, crude protein, carbohydrate and fat content were determined according to AOAC procedures (2002).

### Data analysis

Data were subjected to analysis of variance (ANOVA) in Genstat (11th edition) Statistical Package. Due to their high coefficient of variation (CV), data of ash, crude protein and crude fibre were transformed by square root method. The differences in means were assessed by Fisher’s least significant difference test (LSD) at 5% probability level.

## RESULTS

### Effect of accelerated ageing on seed membrane deterioration

The leachate conductivity of *Tetrapleura tetraptera* seeds increased with increasing ageing and soaking time (Figure 1). The results showed an increase in the conductivity reading of leachate solution of seeds that were subjected to accelerated ageing treatment as compared with the control. It appears that for both traditional and salt saturated methods, longer period of exposure resulted in higher seed leaking. In a similar way, the ageing temperature of 41°C showed greater seed leaking than 38°C. However, the salt saturated accelerated ageing method showed less impact on the conductivity readings of the leachate solutions as compared with the traditional method.

**Figure 1:**
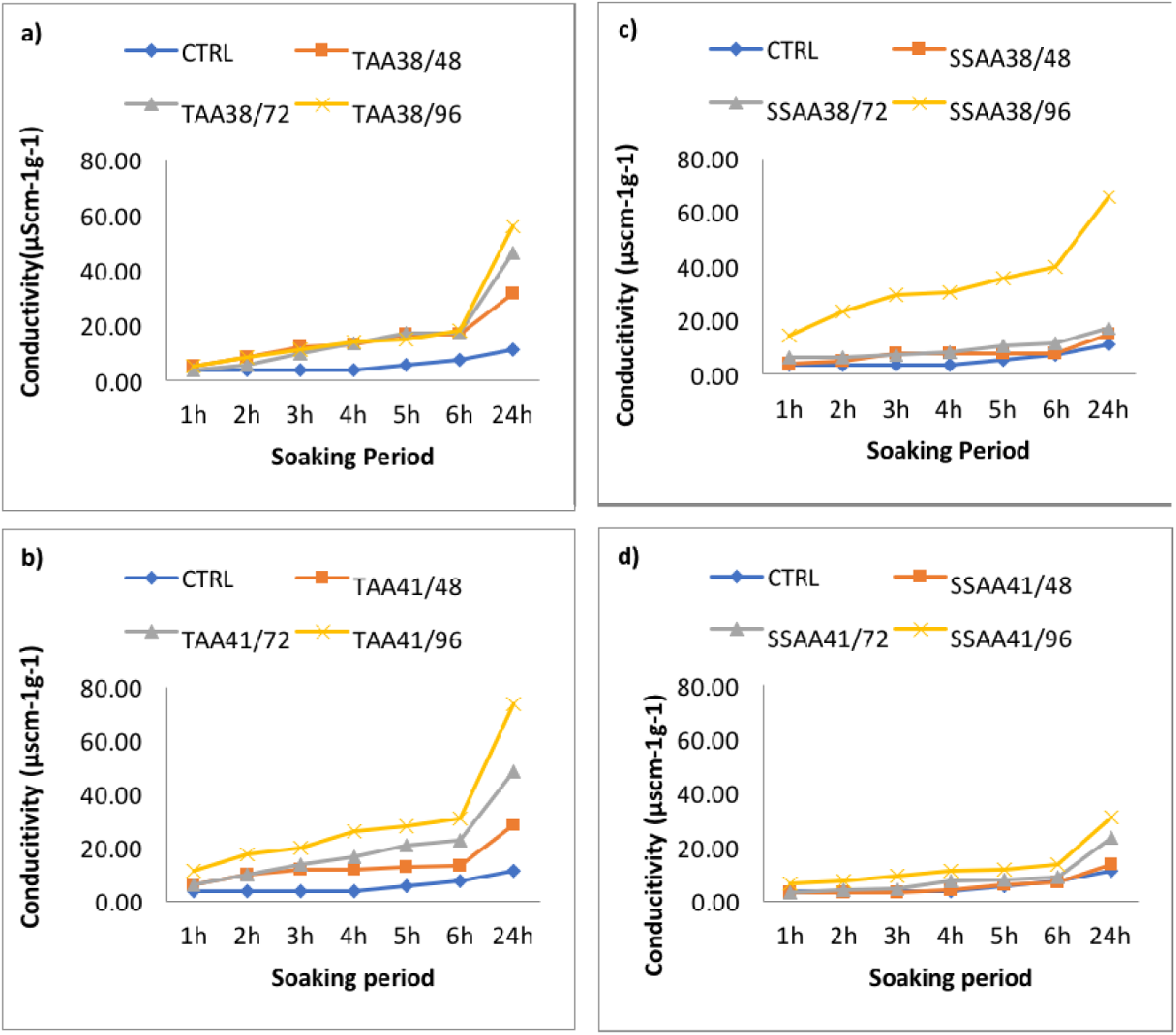
Leachate conductivity of *Tetrapleura tetraptera* seed, as affected by ageing tests **a)** TAA method for 48, 7 and 96 hours at 38°C. **b)** TAA method for 48, 72 and 96 hours at 41°C. **c)** SSAA) method for 48, 72 and 96 hours at 38°C. **d)** SSAA method for 48, 72 and 96 hours at 41°C.

### Effect of accelerated ageing on seed moisture content

The initial seed moisture content was almost identical. The maximum variation among the eight *T tetraptera* seed lots was 0.77% (8.32 to 9.09%). After being exposed to the traditional and salt saturated accelerated ageing methods, *T tetraptera* seeds, showed different moisture absorption patterns as indicated in figure 2. The seed moisture content, after the traditional ageing periods, ranged from 9.12 to 18.37%. Variations in moisture content between seed lots for seeds subjected to traditional accelerated ageing method ranged between 0.6 to 8.16%. For seeds exposed to the salt saturated solution, the moisture content presented smaller and more uniform values ranging from 9.06 to 12.79% for the two temperature regimes after the ageing periods with a moisture variation from 0.27 to 2.04%. Seed moisture content increased after exposure to the various ageing conditions but the traditional accelerated ageing conditions result in large variation in the seed moisture content as compared to the salt saturated conditions. However, the ageing temperature of 38°C showed greater impact on the seed moisture content of the eight lots for both traditional and salt saturated methods.

**Figure 2:**
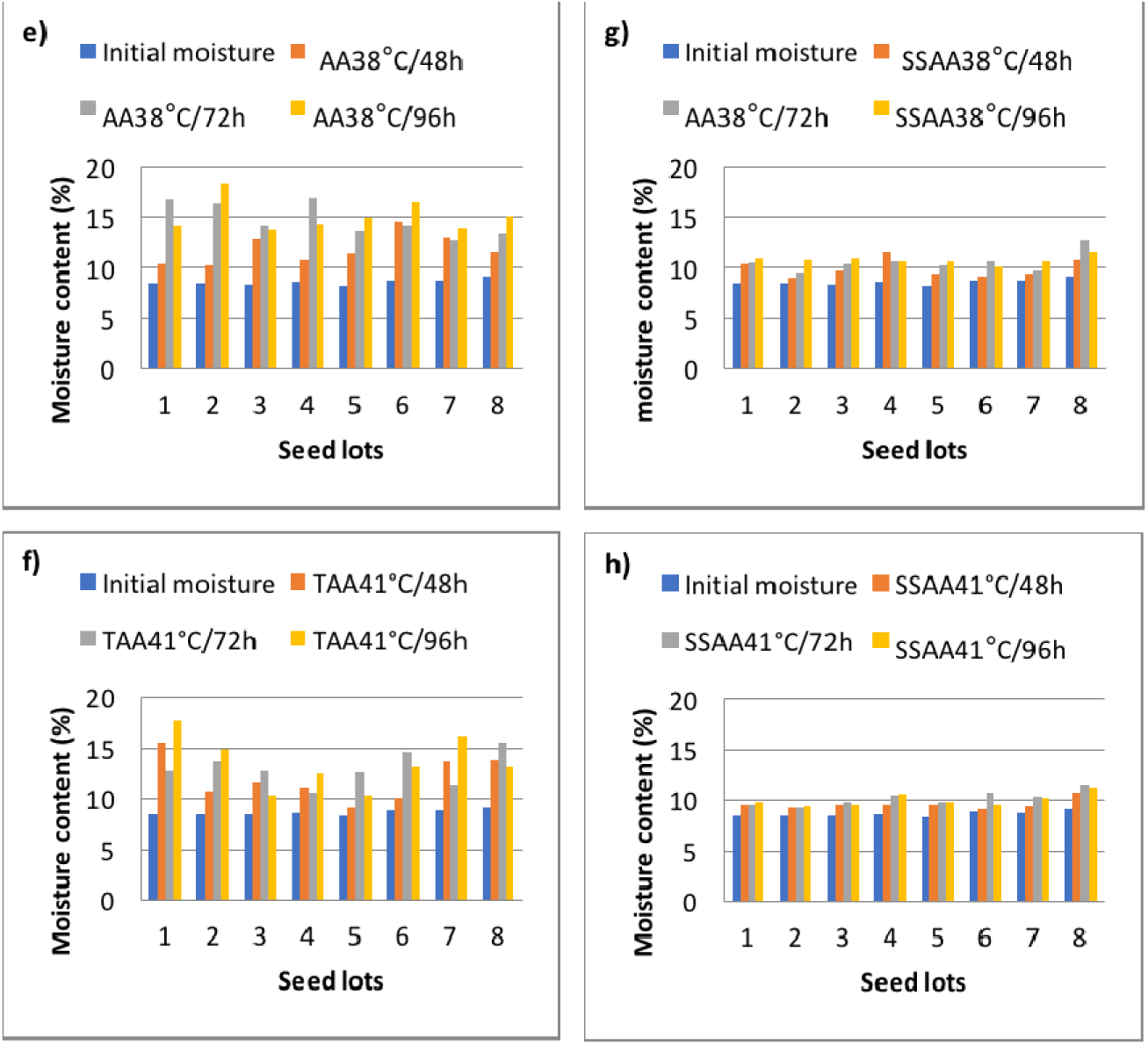
Seed moisture contents variation in *Tetrapleura tetraptera* seed after being exposed to accelerated agein conditions e) TAA method for 48, 72 and 96 hours at 38°C. f) TAA method for 48, 72 and 96 hours at 41°C. g) SSAA) method for 48, 72 and 96 hours at 38°C. h) SSAA method for 48, 72 and 96 hours at 41°C.

### Effect of accelerated ageing conditions on seed chemical composition

#### Ash content (%)

The traditional accelerated ageing method showed no significant differences in the ash content among treatments (Table1). Concerning the salt saturated accelerated ageing method, significant (p<0.01) differences were observed among means for the two temperature regimes (38°C and 41°C), and the interaction temperatures and periods (Table2). However, both traditional and salt saturated ageing methods showed an increase in ash content as compared to the control.

**Table 1:**
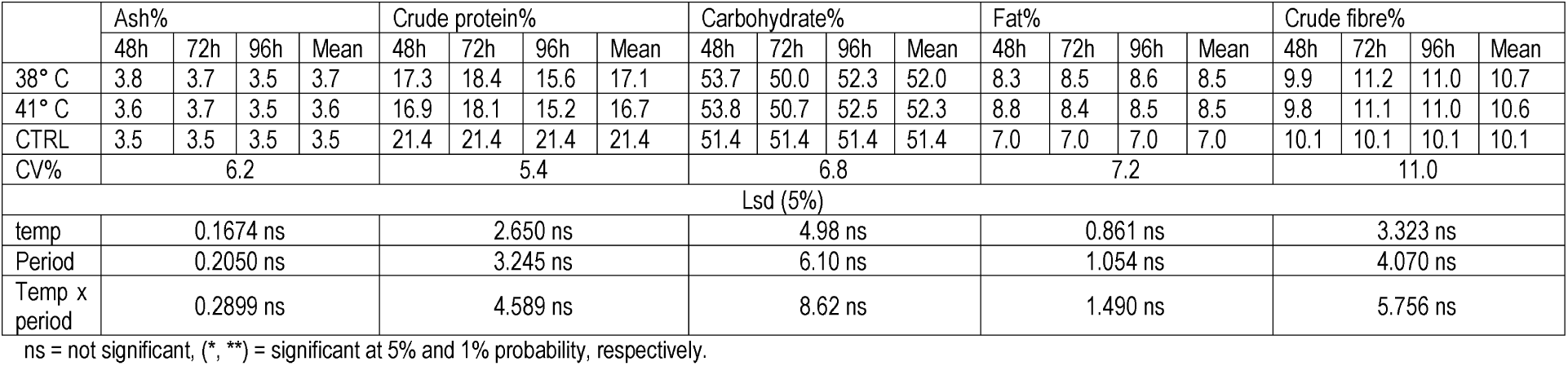
Chemical composition of *Tetrapleura tetraptera* seeds after being subjected to Traditional Accelerated Ageing test for 48, 72 and 96 hours at 38 and 41°C.

**Table 2:** Chemical composition of *Tetrapleura tetraptera* seeds after being subjected to Salt-Saturated Accelerated Ageing test for 48, 72 and 96 hours at 38 and 41°C.

| Tempe<br>ratures | Ash% |  |  |  | Crude protein% |  |  |  | Carbohydrate% |  |  |  | Fat% |  |  |  | Crude fibre% |  |  |  |
| --- | --- | --- | --- | --- | --- | --- | --- | --- | --- | --- | --- | --- | --- | --- | --- | --- | --- | --- | --- | --- |
|  | 48h | 72h | 96h | Mean | 48h | 72h | 96h | Mean | 48h | 72h | 96h | Mean | 48h | 72h | 96h | Mean | 48h | 72h | 96h | Mean |
| 38° C | 3.2 | 3.0 | 3.6 | 3.2 | 18.7 | 19.6 | 14.2 | 17.5 | 53.4 | 53.5 | 55.0 | 54.0 | 7.5 | 6.3 | 7.3 | 7.0 | 11.5 | 10.7 | 12.8 | 11.6 |
| 41° C | 3.8 | 3.8 | 3.4 | 3.7 | 19.3 | 16.2 | 16.1 | 17.2 | 48.7 | 50.7 | 50.7 | 50.1 | 9.3 | 8.3 | 8.5 | 8.7 | 11.1 | 13.4 | 12.3 | 12.3 |
| CTRL | 3.5 | 3.5 | 3.5 | 3.5 | 21.4 | 21.4 | 21.4 | 21.4 | 51.4 | 51.4 | 51.4 | 51.4 | 7.0 | 7.0 | 7.0 | 7.0 | 10.1 | 10.1 | 10.1 | 10.1 |
| CV% | 4.4 |  |  |  | 2.1 |  |  |  | 0.9 |  |  |  | 3.7 |  |  |  | 1.1 |  |  |  |
| Lsd (5%) |  |  |  |  |  |  |  |  |  |  |  |  |  |  |  |  |  |  |  |  |
| Temp | 0.213** |  |  |  | 0.511 ns |  |  |  | 0.697** |  |  |  | 0.408** |  |  |  | 0.193** |  |  |  |
| Period | 0.261 ns |  |  |  | 0.626** |  |  |  | 0.854** |  |  |  | 0.499** |  |  |  | 0.236** |  |  |  |
| Temp<br>x<br>period | 0.369** |  |  |  | 0.885** |  |  |  | 1.208 ns |  |  |  | 0.706 ns |  |  |  | 0.334** |  |  |  |

ns = not significant, (\*, \*\*) = significant at 5% and 1%probability, respectively.

#### Crude protein (%)

The traditional accelerated ageing method had no significant (p<0.01) effect on the crude protein content (Table1), while the salt saturated method showed significant (p<0.01) differences among means (Table2). The mean protein content fell from 21.4% in the control to 17.5% and 17.2% (38 and 41°C respectively) for the salt saturated method and to 17.1% and 16.7% (38 and 41°C respectively) for traditional method. The temperature of 41°C largely reduced the protein content of the seeds as compared to the temperature regime of 38°C.

#### Carbohydrate (%)

The salt saturated method showed significant (p<0.01) differences for the temperatures and period but the interaction of the two factors was not significant (Table 1). The differences in carbohydrate content was not significant (p<0.01) for the traditional method (Table1). However, both traditional and salt saturated methods resulted in an increase in carbohydrate content as compared to the control treatment.

#### Fat (%)

There was no significant (p<0.01) variation between the mean fat content of seeds that were subjected to the traditional method (Table 1). Though there was a significant (p<0.01) difference due to the temperatures and periods alone, the interaction of the two factors produced no significant (p<0.01) effect in the salt saturated method (Table 2). Nonetheless, both traditional and salt saturated methods resulted generally in an increase in the fat content as compared with the control treatment.

#### Crude fibre (%)

The crude fibre content of seeds subjected to the traditional method showed no significant differences (p<0.01) (Table1). As for seeds that were subjected to the salt saturated methods, there was a significant (p<0.01) difference in crude fibre content for the temperatures (38°C and 41°C), the periods and their interactions (Table 2). There was an increase in the crude fibre content with the combination of 41°C and 72h having the highest value (13.4%) and 38°C and 72h having the lowest (10.7%).

## DISCUSSION

The initial seed moisture content was within the limits of two percent points required for the consistency of the ageing results (Marcos-Filho, 1999). The moisture content after the ageing treatments, results were in general similar for the eight lots under study but varied with the ageing methods, temperature and exposure time. The traditional accelerated ageing showed large variations in the seed moisture content (from 0.6 to 8.16% depending on the period and temperature combination), as a result of the high relative humidity (100% RH) to which seeds were subjected. The variation in moisture content of seeds subjected to the traditional accelerated ageing exceeded the tolerable limit for conducting the test (3 to 4%), according to Marcos Filho (1999). However, the moisture content of seeds subjected to the salt saturated conditions showed smaller and more uniform values (0.27 to 2.04% difference), after the ageing periods, as compared to those observed for seeds aged using the traditional method. The observations of Jianhua & McDonald (1996) which stipulated that the use of saturated salt method contributed to slow down of moisture absorption by seeds were then confirmed for *Tetrapleura tetraptera* seed. In addition, the use of salt-saturated solutions had also contributed to reduce or inhibit the growth of fungi, thus reducing the sources of variation for the results (Jianhua & McDonald, 1996).

The loss of membrane integrity is believed to be one of the primary reasons for loss of viability (Malik and Jyoti, 2013). Seed membrane deterioration results in an increase in cell permeability, allowing large quantities of cellular components to diffuse out when the seed is soaked in water (Brasavarajappa et al., 1991). An increase in conductivity of the leachate solution in the accelerated-aged seed as compared to seeds that were not subjected to the ageing treatments was observed. This indicated that accelerated ageing triggered *T tetraptera* seed membrane deterioration and thereby released its seed coat dormancy. According to Chaisurisri et al. (1993), the increase in membrane permeability in the accelerated-aged seed is possibly due to changes that occurred in the molecular structure of the membrane.

The increase in the metabolism during ageing reduced food reserves and, subsequently, seed vigour declined (Blanche et al., 1990). This study clearly revealed changes in chemical composition of *T*etrapleura *tetraptera* seeds exposed to accelerated ageing conditions. When compared to the initial chemical composition of seeds of the species, the seeds subjected to the various accelerated ageing condition showed a large variation in the crude protein, carbohydrate, fat, ash and crude fibre content. In fact, the crude protein content decreased in the aged seeds from 21.4% in the initial seed to a minimum of 14.2% depending on the ageing method, temperature and period. Meanwhile, the carbohydrate, fat, ash and crude fibre content increased significantly in the aged seeds. These results corroborate the work of Verma et al. (2003) that reported an increase in carbohydrate content and a decrease in protein content of deteriorating seeds. In addition, Sossou et al. (2017) reported that the tested ageing conditions led to changes in *T tetraptera* seed germination percentage. Obviously, the loss of membrane integrity and changes in seed composition observed in this study triggered seed coat dormancy release and seed deterioration in *T tetraptera* seed, thereby a decrease in seed viability and physiological quality.

## CONCLUSION

This research showed that the accelerated ageing process have significant effect on membrane integrity and chemical composition of *T*etrapleura *tetraptera* seed. Despite the fact that the different tested conditions were less efficient in segregating *T tetraptera* seed lots into different vigour level, the accelerated ageing process triggered seed coat dormancy release and seed deterioration in *T tetraptera* seed. The accelerated ageing test combined with a storage experiment could therefore be a promising technique for estimating the storage potential of *T tetraptera* seed. However, further investigations are required to assess other conditions (temperature and time of exposure combination) taking into account the enzyme activity to yield better understanding and standardize the appropriate testing conditions for the species.

## ACKNOWLEDGEMENT

The authors are grateful to the CSAA Intra-ACP mobility project and the Regional Universities Forum for Capacity Building in Agriculture (RUFORUM) for supporting this work. This paper is a contribution to the 2018 Sixth African Higher Education Week and RUFORUM Biennial Conference.

